# Expression of a *Malassezia* codon optimized mCherry fluorescent protein in a bicistronic vector

**DOI:** 10.1101/2020.04.16.044297

**Authors:** Joleen P.Z. Goh, Giuseppe Ianiri, Joseph Heitman, Thomas L. Dawson

## Abstract

The use of fluorescent proteins allows a multitude of approaches from live imaging and fixed cells to labelling of whole organisms, making it a foundation of diverse experiments. Tagging a protein of interest or specific cell type allows visualization and studies of cell localization, cellular dynamics, physiology, and structural characteristics. In specific instances fluorescent fusion proteins may not be properly functional as a result of structural changes that hinder protein function, or when overexpressed may be cytotoxic and disrupt normal biological processes. In our study, we describe application of a bicistronic vector incorporating a Picornavirus 2A peptide sequence between a NAT antibiotic selection marker and mCherry. This allows expression of multiple genes from a single open reading frame and production of discrete protein products through a cleavage event within the 2A peptide. We demonstrate integration of this bicistronic vector into a model *Malassezia* species, the haploid strain *M. furfur* CBS 14141, with both active selection, high fluorescence, and proven proteolytic cleavage. Potential applications of this technology can include protein functional studies, *Malassezia* cellular localization, and co-expression of genes required for targeted mutagenesis.

## Introduction

*Malassezia*, comprising 18 currently recognized species, are a unique group of lipophilic basidiomycetes that evolved independently from related plant pathogen lineages (Xu et al. 2007), and represent a ubiquitous and dominant eukaryotic microbial community on human skin (Grice and Segre 2011; Findley et al. 2013; Grice and Dawson 2017). In adaption to life on mammalian skin, *Malassezia* genomes have been reshaped to secrete an armory of proteases, lipases, and phospholipases, amongst other enzymes to support their growth. These skin-dwelling yeasts usually maintain a symbiotic relationship with their human host but can quickly shift into opportunistic pathogens, usually induced by environmental alterations such as breaches in skin barrier integrity, dysfunctional immune response, or age related changes that affect skin, such as aging, puberty, or menopause (Grice and Segre 2011). *Malassezia* are the causative agents of dandruff and seborrheic dermatitis, and are associated with myriad clinical conditions such as atopic dermatitis, pityriasis versicolor, and folliculitis (Gaitanis et al. 2012; Theelen et al. 2018). Beyond superficial cutaneous disorders, *Malassezia* are also responsible for catheter-associated infections and invasive septic fungemia (Barber et al. 1993; Gaitanis et al. 2012; Kaneko et al. 2012; Iatta et al. 2014). Recent reports illustrate pathogenic roles for *Malassezia* in Crohn’s Disease and pancreatic ductal adenocarcinoma through an elevated inflammatory response linked to CARD-9 and the complement cascade (Limon et al. 2019; Aykut et al. 2019). Relatively little is known about the specific mechanisms of *Malassezia* pathogenesis, despite decades of investigation and their broad significance, meaning much remains to be explored about this important group of yeasts. Research efforts on *Malassezia* are gaining traction as more species are being identified in human gut microflora, animal skin, and even in Antarctic and marine environments, making them amongst the most ubiquitous of fungi (Amend 2014; Theelen et al. 2018). These new findings have rapidly increased interest, and are driving advances in diverse species identification, definition of axenic culture conditions, and development and application of tools to dissect *Malassezia* genomic complexity (Dawson 2019).

*Malassezia* were widely accepted as highly recalcitrant to conventional transformation techniques including biolistic, electroporation, and lithium acetate methods. Recent developments in genetic modifications with the use of *Agrobacterium* have not only established gene transfer, but also provided tremendous headway in studies of previously uncharacterized *Malassezia* gene function (Ianiri et al. 2016; 2019; Sankaranarayanan et al. 2020). Earlier studies have also applied the use of fluorescence protein tagging (Celis et al. 2017; Sankaranarayanan et al. 2020). Additionally, the simultaneous co-expression of multiple genes and fluorescent proteins has applications ranging from monitoring gene expression (Rasala et al. 2012; Lewis et al. 2015), protein tagging, to live cell or whole organism labelling and imaging (Provost, Rhee, and Leach 2007; Kim et al. 2011; Ahier and Jarriault 2014), enabling this technique as a cornerstone in biomedical research.

One of the most common approaches in collective gene expression exploits the incorporation of the 2A oligopeptide, first identified in viral genome of foot-and-mouth disease virus (F2A). Subsequently, other 2A sequences were discovered in porcine teschovirus-1 (P2A), equine rhinitis A (E2A), and *Thosea asigna* virus (F2A). 2A peptides are usually between 18-22 residues and reports have suggested 2A sequences encode a single open reading frame (ORF), and impedes the formation of a peptide bond between glycine and proline residues, allowing the generation of discrete protein products (Trichas, Begbie, and Srinivas 2008; Lewis et al. 2015). The use of 2A sequence overcomes the need of bidirectional or multiple promoters and skips use of numerous co-transfection plasmids, offering greater efficiency and simplicity in construct designs and transformation.

In this report, we demonstrated use of a bicistronic expression system to simultaneously produce active fluorescent mCherry and dominant nourseothricin resistance (NAT) non-fusion proteins, delivered via *Agrobacterium tumefaciens*-mediated transformation (ATMT) in *Malassezia furfur*. This is achieved through incorporating viral P2A sequence between gene ORFs, with mCherry upstream of NAT, designed to ensure expression of mCherry if NAT protein is expressed. *Agrobacterium* Transfer DNA (T-DNA) expression vector was designed and constructed with unique restriction enzyme sites to allow straightforward modification to any of the genetic elements to support broad experimental use and complement alternative experimental needs.

## Results

### Random Insertional Mutagenesis in *M. furfur*

The expression vector was first constructed in *E. coli* pUC57 cloning vector by insertion between the actin *ACT1* promoter and terminator of *Malassezia sympodialis* ATCC 42132, a codon optimized mCherry gene and a P2A viral sequence (Supplementary Table 1 and Supplementary Figure 1) followed by the *NAT* selection marker gene, forming a single ORF. The expression cassette was digested from purified pUC57 and cloned into an *Agrobacterium* tumor-inducing (Ti) backbone binary vector, generating the resulting plasmid, pJG201702 (Figure 1). The bicistronic expression plasmid pJG201702 was verified by PCR and Sanger sequencing (data not shown) prior to electroporation into competent *Agrobacterium tumefaciens*.

**Figure 1.**
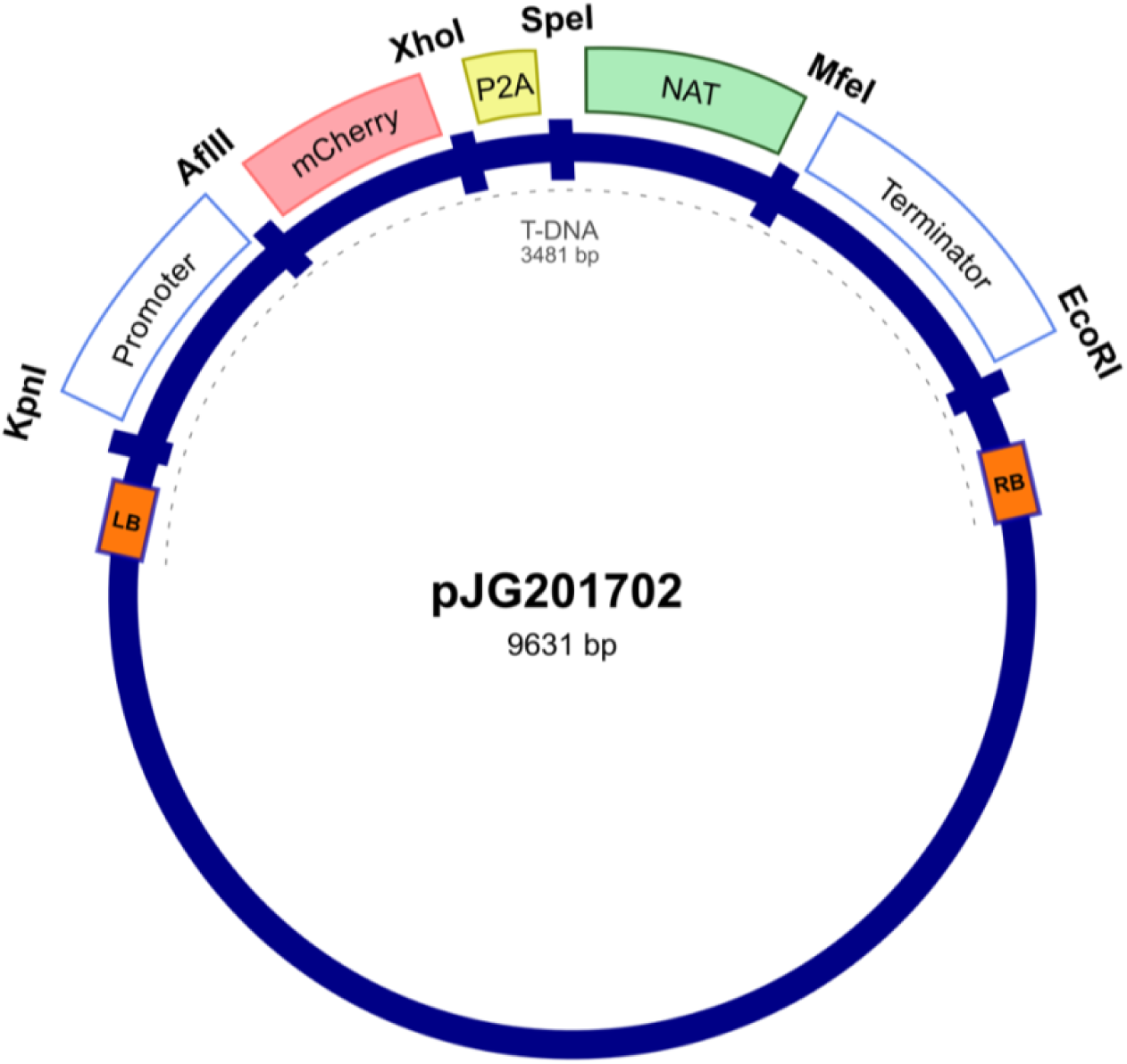
Random integration bicistronic vector. Generation of bicistronic construct comprised use of promoter and terminator sequences from *Malassezia sympodialis* ATCC 42132 in *Agrobacterium tumefaciens* backbone vector. Actin encoding regulatory elements govern the sequential expression of mCherry and nourseothricin sulfate (NAT) antibiotic resistance marker, separated by porcine teschovirus-1 2A (P2A) pseudo-autolytic cleavage sequence. pJG201702 vector was created with multiple restriction cut sites between each genetic element for future modification.

Transformations were carried out as previously described (Ianiri et al. 2016; Celis et al. 2017), and randomly selected NAT-resistant colonies were analyzed by PCR to detect the presence of *NAT* and mCherry ORFs. Predicted amplicons of 576 bp and 708 bp for *NAT* and mCherry respectively, were observed in 4 out of 23 transformants but absent in wild type (Figure 2A). The recovery of 17 false positive NAT-resistant colonies are likely to be attributed to spontaneous mutations and this chemically induced resistance to NAT was also observed in previous work (Ianiri et al. 2016). The 4 mCherry- and NAT-positive transformants and wild type were serially diluted in PBS and spotted on mDixon and selective medium-supplemented with NAT. Wild type CBS14141 was not able to proliferate in the presence of NAT while engineered *M. furfur* strains displayed growth similar to wild type on mDixon (Figure 2B). These results suggest in the selected transformants, the *NAT* resistance gene was incorporated into the genome, expressed, and translated into a functional protein without affecting fitness.

**Figure 2.**
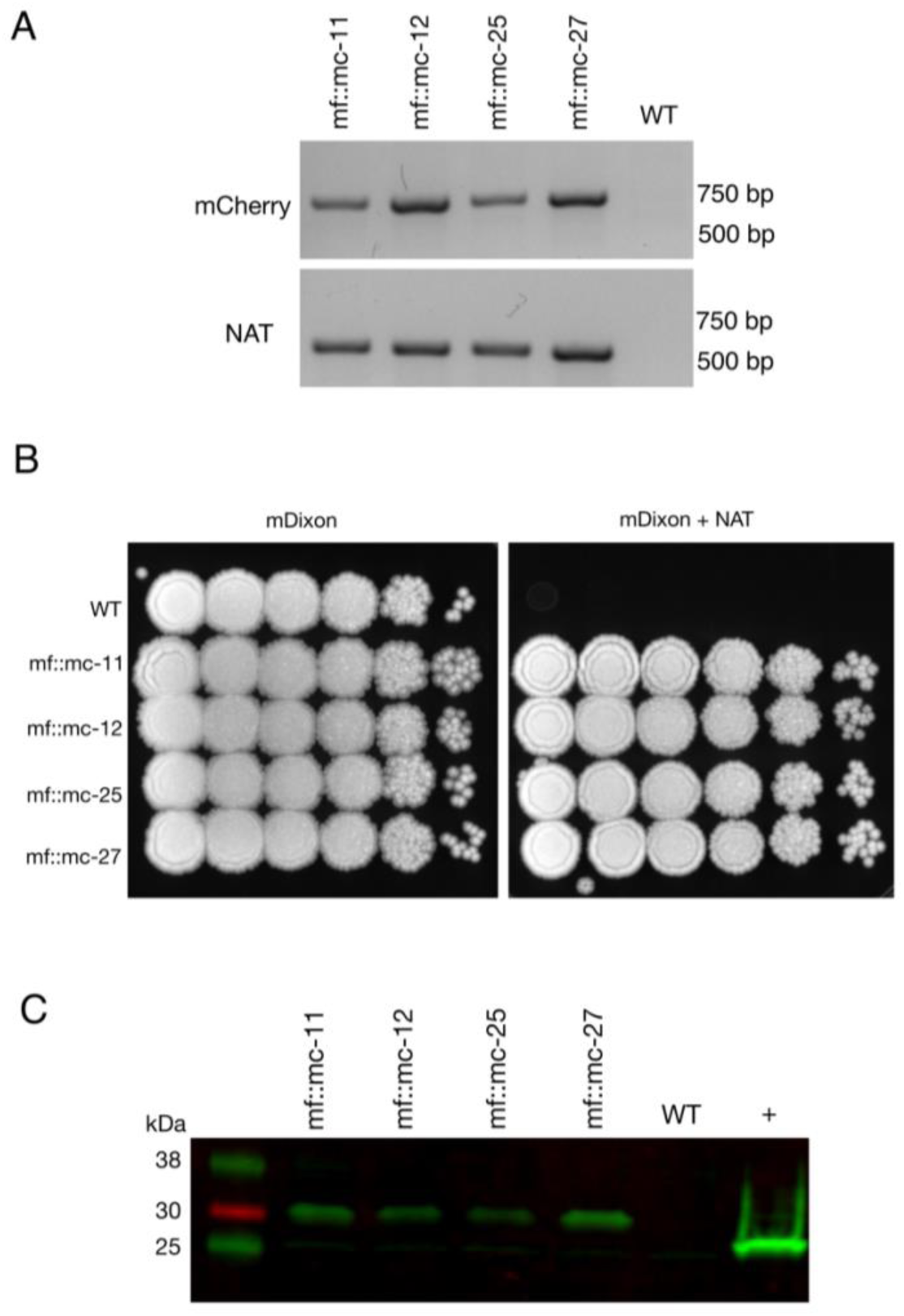
Characterization and analysis of transformants. **(A)** PCR screening of transformants and wild-type cells were analyzed with mCherry and NAT ORF spanning primers, indicating the appropriate sized product in transformants and not wild-type cells. **(B)** Transformants and *M. furfur* CBS 14141 wild-type clones were ten-fold serially diluted in PBS and 3uL of cell suspensions were spotted on mDixon growth medium and selection media-supplemented with 100ug/mL NAT, indicating NAT resistance in selected clones. **(C)** Immunoblot was performed with whole cell lysates, probed with anti-RFP primary antibodies. Transformants expressing mCherry contained additional amino acid residues from viral P2A protein tag, produced expected protein bands with an upward shift of molecular weight at 29 kDa. In comparison, 26.7 kDa native molecular weight band was detected in positive control, TurboRFP expressing keratinocyte cell lysate.

### P2A cleavage efficiency

To determine whether P2A peptide allows proper and efficient cleavage and release of the mCherry and NAT proteins, total protein lysates were prepared and analyzed from wild type and transformants. Immunoblot analysis using a red fluorescent protein antibody (RFP, mCherry) revealed the expected single band of 29 kDa with no observable band detected in wild type lysate (Figure 2C).

### *In vivo* fluorescence assessment

*M. furfur* CBS 14141 wild type and three selected genetically engineered strains, mf::mc-12, -25, and -27 were fluorescently imaged. Live cell imaging revealed transformants exhibit a higher fluorescence signal compared to wild type (Figure 3A). Wild type displayed classical autofluorescence observed in *Malassezia* species that localizes mainly in the yeast cell wall, while all selected transformants displayed a strong mCherry cytoplasmic signal, with mf::mc-27 demonstrating the strongest fluorescence. In an effort to reduce the autofluorescence background, mf::mc-27 and wildtype were cultured in liquid media and cells were collected at mid-log phase to avoid the accumulation of fluorescent metabolites, dead or stationary cells. Further, cells were washed in PBS to remove residual culture media before imaging but detectable autofluorescence was still observed in wildtype. Regardless, mf::mc-27 transformant demonstrated higher fluorescence, an indicator that the mCherry protein was expressed and functional (Figure 3B).

**Figure 3.**
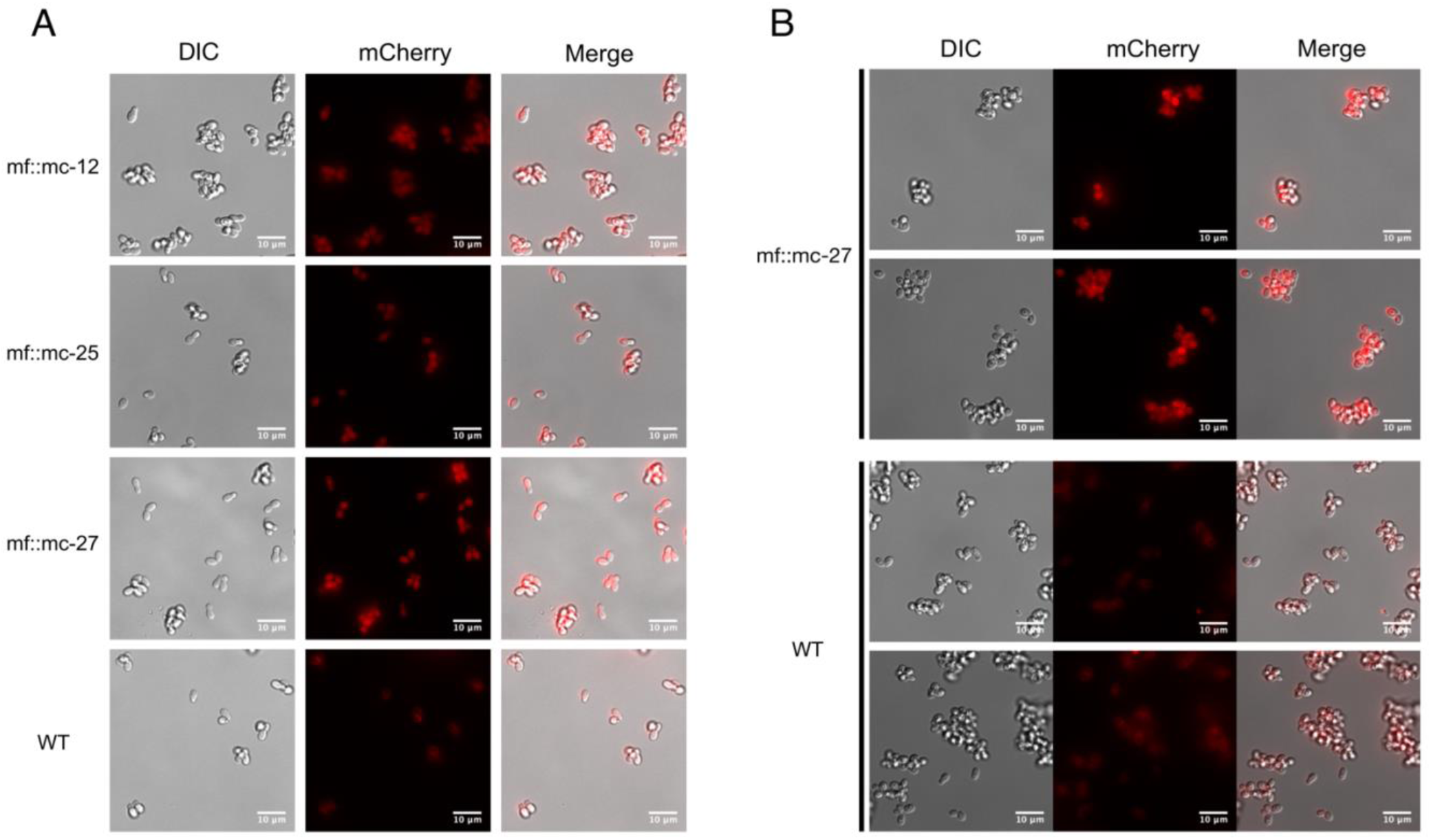
Live fluorescent Imaging. **(A)** Transformants and wild-type cells were collected from mDixon agar and imaged under TRITC channel and differential interference contrast (DIC). Wild-type cells displayed localized autofluorescence in cell wall, in contrast to transformants exhibiting stronger fluorescence distributed uniformly throughout the cells. **(B)** Cells in exponential growth phase were collected to minimize accretion of dead or stationery cells and aggregation of fluorescence metabolites. Transformant mf::mc-27 presented higher fluorescence intensity in comparison to *M. furfur* CBS 14141 wild type cells which displayed persistent low levels of fluorescence background.

### Location of the inserted genetic cassette

Transformant mf::mc-27 was subjected to Illumina sequencing (Novogene) to determine the site of the T-DNA insertion. Sequenced DNA fragments were mapped to *M. furfur* CBS 14141 reference genome assembly (Sankaranarayanan et al. 2020) and RNAseq data (TLD lab, unpublished data). Bioinformatic analysis located the insertion of the exogenous DNA cassette in an intergenic region on chromosome 2 (Figure 4). Sanger sequencing of T-DNA through to upstream chromosomal DNA confirmed the presence of a complete ORF of a putative CDC25-related phosphatase gene. PCR amplifications of the downstream chromosomal DNA from the insertional point were unsuccessful. However, Illumina sequencing reads were able to detect both intact transformation DNA cassette and predicted ORF of the adjoining adiponectin receptor.

**Figure 4.**
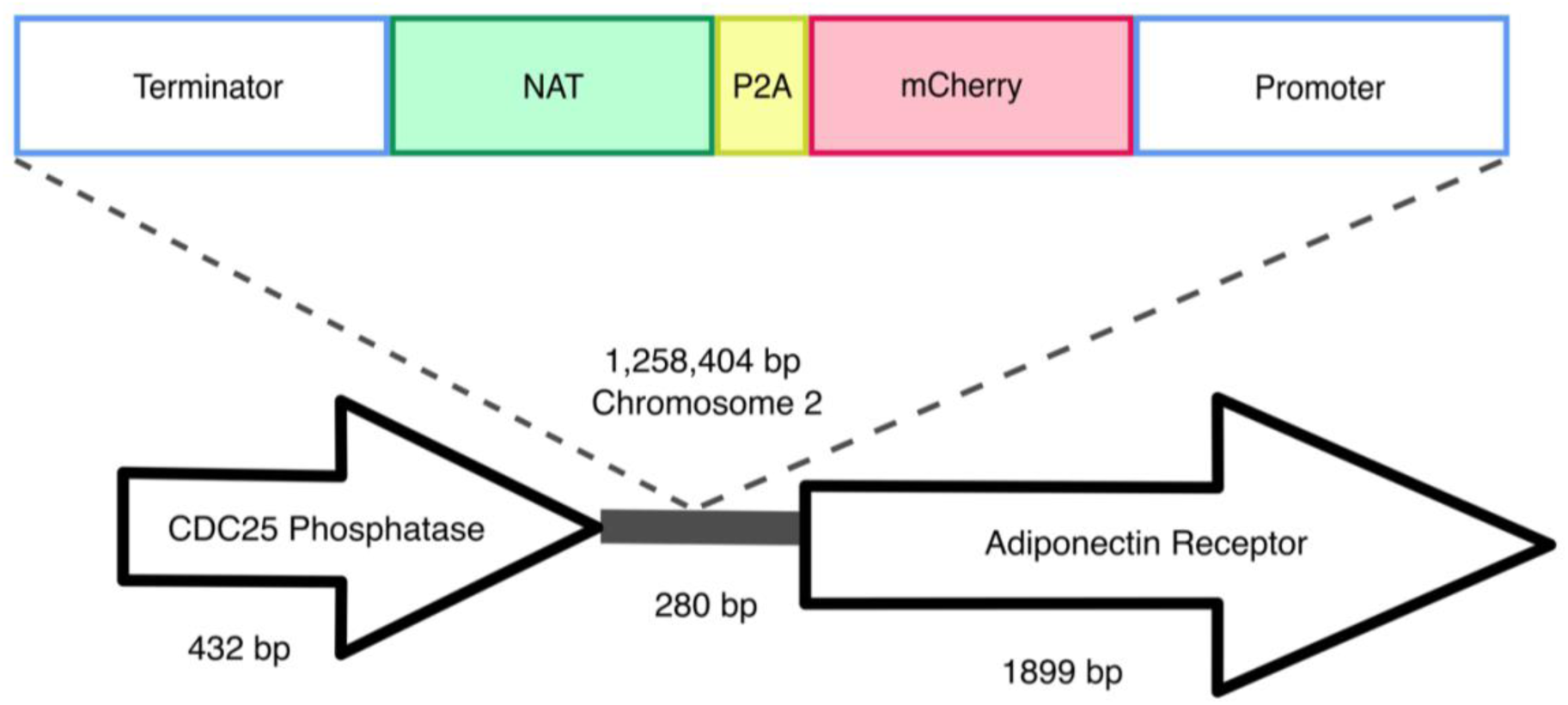
Random insertional mutagenesis. Using Illumina and Sanger sequencing, exogenous T-DNA was identified in chromosome 2 of mf::mc-27 mutant. Sanger sequencing further validates the presence of CDC25 phosphatase gene, adjacent to exogenous DNA of actin terminator and NAT. Illumina reads were able to detect both complete flanking ORFs of the hypothetical adiponectin rector protein, and exogenous T-DNA containing *Malassezia* actin promoter, terminator, and mCherry-P2A-NAT cassette.

## Discussion

*Malassezia* are indispensable members of a healthy human skin microbiome, found on almost all warm-blooded animals, and can even be traced in a marine ecosystem (Amend 2014; Theelen et al. 2018). This diverse group of lipophilic yeasts is often implicated in various cutaneous diseases and recent studies have identified *Malassezia* as playing pivotal roles in the progression of Crohn’s Disease and exocrine pancreatic cancer (Ashbee and Evans 2002; Limon et al. 2019; Aykut et al. 2019). Yet much of the specific mechanisms and disease pathogenesis remain elusive due to the lack of capability to perform gene studies, hampering the advance of research developments. Today, two research groups have demonstrated the use of a soil bacterium, *Agrobacterium tumefaciens* to genetically modify *Malassezia*, paving a new avenue to investigate functional genomics (Ianiri et al. 2016; Celis et al. 2017). In this study, we leveraged the application of *Agrobacterium-*mediated transformation to introduce a bicistronic expression vector to co-express red fluorescence protein, mCherry, and NAT resistance marker in *Malassezia furfur*.

The usage of inter-kingdom *Agrobacterium*-mediated transformation in delivering exogenous DNA is a convenient and versatile tool applied in diverse engineered eukaryotic species. This pathogenic bacterium effectively housebreaks into the host genome and influences host cellular processes to its benefit. It also provides an inexpensive and flexible approach in designing and utilizing expression vectors. ATMT approach also has its set of limitations including the unpredictable efficiency of transgene expression due to position effects, leading to varying transgenes expression levels among the same pool of transformants. In our study, *Agrobacterium tumefaciens* Transfer DNA (T-DNA) does not contain homologous DNA sequences to *Malassezia furfur* genome, in a deliberate attempt to assess random integration and gene disruption.

The construction of the bicistronic vector, pJG201702 included the use of actin encoding promoter and terminator to regulate the expression of mCherry and NAT proteins. The arrangement of the transcribed proteins was specifically designed to direct the obligatory expression of mCherry when NAT protein is simultaneously expressed. It was engineered to avoid the possibility of selective gene expression as *Malassezia* acquired the potential to undergo genomic rearrangements, excising non-essential genes in maintaining a condensed genome (Xu et al. 2007; Sankaranarayanan et al. 2020).

Generation of non-fusion proteins was facilitated through the insertion of P2A viral sequence between mCherry and NAT. Ribosomal skipping of the peptide bond formation between glycine and proline residues within P2A sequence results in a pseudo-cleavage event to take place, separating the two protein products (Kim et al. 2011; Szymczak-Workman, Vignali, and Vignali 2012; Liu et al. 2017). In pJG201702, each genetic component is flanked by restriction enzyme sites, designed for the ease of future modification including the development of multi-cistronic vectors through incorporating multiple 2A sequences to generate varied gene products. Additionally, the mCherry and P2A sequences were codon optimized to improve protein expression in *Malassezia*.

To assess the functionality of P2A sequence in the generation of discrete mCherry and NAT protein products, a monoclonal antibody specific for red fluorescent protein was probed against whole cell lysates extracted from wild type and transformants. We detected the predicted protein band at 29 kDa in all transformants. The pseudo autolytic-cleavage occurs near the end of P2A sequence resulting in the retention of a 2A tag (21 amino acid residues) at the end of mCherry C-terminus, explaining the significant shift of the cleaved mCherry protein from its native weight of 26.7 kDa. This suggests adequate self-cleaving efficiency with the inclusion of a Gly-Ser-Gly (GSG) linker in P2A residues which promotes cleavage efficiency (Wang et al. 2015; Kim et al. 2011). Correct functionality of the generated vector was demonstrated by strong mCherry expression of NAT resistant transformants compared to wild type, which displayed autofluorescence background probably due to the presence of lipids within the growth media (Croce and Bottiroli 2014) and the production of naturally occurring intracellular metabolites (Mayser et al. 2002; Maslanka, Kwolek-Mirek, and Zadrag-Tecza 2018).

We sequenced the genome of one fluorescent transformant and confirmed correct T-DNA integration in *M. furfur* genome. The T-DNA was inserted between two genes, the gene upstream of T-DNA encodes a putative CDC25-related phosphatase, an yeast ortholog of a Ras guanyl-nucleotide exchange factor. This gene is thought to be involved in Ras protein signal transduction and cell cycle regulation, traversing the start control point of the mitotic cell cycle (Chen et al. 2000). Downstream of the T-DNA insertion encodes for an adiponectin receptor gene that is predicted to be associated with zinc ion homeostasis (Lyons et al. 2004). Based on the orientation of the genes, it is likely that the T-DNA integrated in the terminator region of CDC25-related phosphatase and the promoter region of the predicted adjoining adiponectin-encoding gene. Intergenic insertions of T-DNA are very common and represent a disadvantage of ATMT in fungi, and they are probably due to a preference for the T-DNA to insert in low-transcribed regions (Michielse et al. 2005; Idnurm et al. 2017). Intergenic insertions were also reported in two previous studies in *Malassezia*, suggesting that additional mechanisms that favor insertion in a ORF-free regions may exist (Ianiri et al. 2016; 2019); this is surprising if we consider that *Malassezia* genomes are consistently small and compact (Wu et al. 2015). This intergenic insertion may explain the lack of phenotypic abnormalities or variations in fitness in mc::mf-17 transformant. Intergenic insertions can also be turned into an advantage by using a conditional promoter that drives the expression of the gene downstream the T-DNA insertion (Kilaru et al. 2015; Ianiri, Boyce, and Idnurm 2017). Another advantage could be the use of non-protein coding region as a safe haven for future genetic engineering in *Malassezia*, especially useful for the reintroduction of genes as complementation without compromising cell viability or disruption of neighboring genes as reported in *C. neoformans* (Arras et al. 2015; Upadhya et al. 2017).

We demonstrated the effective use of a bicistronic vector in *Malassezia*, to generate a codon optimized RFP in a NAT-resistant strain. The application of a bicistronic system can potentially be expanded to a multi-cistronic construct with the aid of multiple cloning sites, to include the expression of assorted genes according to the users’ preference. Most of current research models in understanding skin health include the use of transgenic mice expressing green fluorescence protein (GFP), mammalian cell culture, reconstructed skin epidermis (RHE) or even *ex-vivo* skin models, which can be complemented with these fluorescently labeled *Malassezia*. In addition, the competence to fluorescently label yeast cells, overexpression of genes, and targeted gene replacement can be applied to deepen our understanding of *Malassezia* gene functions and host-microbes interactions.

## Methods

### Strains and culture conditions

Haploid strain *M. furfur* CBS 14141 was obtained from Westerdijk Fungal Biodiversity Institute. Cells were cultured at 32°C, with mDixon medium (36 g/L malt extract, 20 g/L desiccated oxbile, 6 g/L peptone, 2 ml/L glycerol, 2 ml/L oleic acid, 10 ml/L Tween 40, pH 6). Genetically transformed cells were maintained on mDixon supplemented with 100 µg/ml nourseothricin sulfate (NAT).

### Plasmid and constructs

pJG201701 was first constructed with *M. sympodialis* ATCC 42132 actin promoter and terminator sequences, encoding for codon-optimized fluorescent mCherry and porcine teschovirus-1 2A (P2A) sequences (as detailed in Supplementary Table 1), followed by NAT antibiotic resistance marker (GenScript) in pUC57 cloning vector. To enable the translation of both proteins, the stop codon of mCherry was removed and retaining a single stop codon in the second protein, NAT. Both actin regulatory and NAT sequences were taken from pAIM2 (Ianiri et al. 2016). Each genetic elements are adjoined with restriction cut sites to allow efficient modification of the construct. pJG201701 was digested with EcoRI and KpnI, and gene fragments were cloned in pPZP201-BK (Covert et al. 2001), which contains *Agrobacterium tumefaciens* backbone vector using T4 DNA ligase (New England Biolabs). Resultant plasmid, pJG201702 was transferred to *A. tumefaciens* EHA105 strain via electroporation and verified by digestion pattern and PCR.

### *A. tumefaciens*-mediated transformation

Transformation of *M. furfur* were performed using previously established protocol (Celis et al. 2017; Ianiri et al. 2016). *A. tumefaciens* harboring pJG201702 was grown overnight in Luria-Bertani medium supplemented with 50 µg/ml kanamycin at 30°C and 250 rpm in a shaking incubator. Aliquot of cells were used as inoculum, resuspended in induction medium (IM) containing 100 µM acetosyringone (Sigma) and cultured for additional 6 hours to OD_600_ of 1. *M. furfur* CBS 14141 cells were collected at mid-log phase and used at OD_600_ of 1. Proportional volumes of bacterial and yeast cells were mixed and filtered through 0.45 µm mixed cellulose membrane (Merck, Millipore) before transferring onto IM agar supplemented with 200 µM acetosyringone. Co-culture plates were incubated at 25°C for 5 days, after which cells were washed in 20 mL sterile PBS and plated on selection medium – mDixon agar containing 100 µg/ml NAT to select for transformants, and 200 µg/ml cefotaxime with 10 µg/ml tetracycline to halt the growth of *A. tumefaciens*.

### Molecular analysis

Genomic DNA of wild-type and engineered strains of *M. furfur* CBS 14141 were isolated with MasterPure yeast DNA kit (Epicentre) as per manufacturer’s protocol with additional step of homogenizing at 6 m/s for 50 sec (MP Biomedicals). PCR amplification of mCherry and NAT genes with mCherry-forward (5’-ATGGTGTCGAAGGGCGAG-3’) and mCherry-reverse (5’-CTTGTAGAGCTCGTCCATGC-3’), and NAT-F (5’-ATGGCGGCCGCCACTCTTGAC-3’) and NAT-R (5’-TTATGGACAAGGCATACTCATATAAAG-3’) primers respectively to screen for positively transformed *Malassezia*.

### Total protein extraction and immunoblot

Washed transformant and wild type yeast cells were collected by centrifugation and resuspended in lysis buffer (50mM Tris-HCl, pH 7.5) with added protease inhibitor cocktail (Nacalai tesque). Cells were mechanically lysed with mixture of ceramic and glass beads (Lysing Matrix E, MP Biomedicals) for 50 seconds and repeated twice to achieve complete cell breakage. Cell debris were separated by centrifugation at 12,000 rpm at 4°C for 5 min and supernatants were transferred to new tubes for repeated centrifugations. Prepared protein samples were separated using 4-20% Tris-glycine SDS-PAGE gradient gel and blotted to PVDF membrane. The blot was blocked in Intercept PBS blocking buffer (Li-cor) and subsequently incubated with RFP monoclonal antibody (ThermoFisher, MA5-15257) and probed with fluorescent anti-mouse secondary antibody. Protein bands were visualized with Odyssey CLx digital imaging system (Li-cor).

### Genome sequencing

Mf::mc-27 isolate was selected for whole genome analysis and nucleic acid was extracted using method described previously. Whole genome sequencing was performed with Illumina Novoseq 6000 at 100x coverage, from 150 bp short insert paired-end reads (NovogeneAIT). CDS prediction using Augustus gene prediction tool identified putative CDC25 Phosphatase and Adiponectin Protein Receptor proteins (Supplementary Table 2).

### Fluorescence microscopy

Live cell imaging was obtained on inverted wide-field microscope (Olympus IX-83). Fluorescence and differential interference contrast (DIC) images were achieved with 60x 1.2 oil objective (plan-Apochromat). Exposure time was kept at 700 ms, and cells were held at 37 °C with applied CO_2_ incubator chamber during image acquisition. Images were processed and analyzed with FIJI (ImageJ).

## Acknowledgements

This study was supported by Agency for Science, Technology and Research IAF-PP (HBMS) grant Asian Skin Microbiome Project H18/01a0/016 (to TLD). Work in the Heitman lab was supported by NIH/NIAID R37 grant AI39115-22 and NIH/NIAID R01 grant AI50113-15. Prof Joseph Heitman is the co-director and fellow of the CIFAR program Fungal Kingdom: Threats & Opportunities. We would like to thank Dr Andrea Camattari for providing the *Malassezia sympodialis* codon usage table, Dr Teun Boekhout at the Westerdijk Fungal Biodiversity Institute for providing the *Malassezia* strains, and Dr Simon L.I.J. Denil for his help with bioinformatic analysis.

## Author Contributions Statement

JPZG, GI, JH and TLD designed the experiments. JPZG performed the experiments and data analysis. JPZG and TLD wrote the manuscript.

