## Supplementary Material for "Expression of a *Malassezia* codon optimized mCherry fluorescent protein in a bicistronic vector"

| Amino Acid | Codon | Frequency % |
|---|---|---|
| * | TAA | 29.39 |
| | TAG | 56.46 |
| | TGA | 14.16 |
| A | GCA | 16.00 |
| | GCC | 27.63 |
| | GCG | 40.98 |
| | GCT | 15.39 |
| C | TGC | 64.41 |
| | TGT | 35.59 |
| D | GAC | 61.90 |
| | GAT | 38.10 |
| E | GAA | 28.24 |
| | GAG | 71.76 |
| F | TTC | 52.82 |
| | TTT | 47.18 |
| G | GGA | 11.61 |
| | GGC | 51.64 |
| | GGG | 13.02 |
| | GGT | 23.74 |
| H | CAC | 59.96 |
| | CAT | 40.04 |
| I | ATA | 7.97 |
| | ATC | 50.54 |
| | ATT | 41.49 |
| K | AAA | 25.79 |
| | AAG | 74.21 |
| L | CTA | 7.22 |
| | CTC | 28.18 |
| | CTG | 35.85 |
| | CTT | 13.57 |
| | TTA | 3.09 |
| | TTG | 12.10 |

| Amino Acid | Codon | Frequency % |
|---|---|---|
| M | ATG | 100.00 |
| N | AAC | 59.38 |
| | AAT | 40.62 |
| P | CCA | 15.36 |
| | CCC | 26.30 |
| | CCG | 36.54 |
| | CCT | 21.79 |
| Q | CAA | 26.10 |
| | CAG | 73.90 |
| R | AGA | 2.31 |
| | AGG | 3.38 |
| | CGA | 8.60 |
| | CGC | 50.29 |
| | CGG | 15.97 |
| | CGT | 19.44 |
| S | AGC | 21.37 |
| | AGT | 11.16 |
| | TCA | 9.77 |
| | TCC | 12.96 |
| | TCG | 31.64 |
| | TCT | 13.10 |
| T | ACA | 19.53 |
| | ACC | 22.05 |
| | ACG | 44.98 |
| | ACT | 13.44 |
| V | GTA | 9.63 |
| | GTC | 27.61 |
| | GTG | 50.99 |
| | GTT | 11.76 |
| W | TGG | 100.00 |
| Y | TAC | 64.59 |
| | TAT | 35.41 |

**Supplementary Table 1**. Relative synonymous codon usage in *Malassezia sympodialis*.

**mCherry**

ATGGTGTCGAAGGGCGAGGAAGACAACATGGCGATCATTAAGGAGTTCATGCGCTTTA
AGGTGCACATGGAGGGCTCGGTGAACGGCCACGAGTTCGAGATTGAGGGCGAGGGC
GAGGGCCGCCCTTACGAGGGCACGCAGACGGCGAAGCTCAAGGTGACGAAGGGCG
GCCCCCTGCCCTTTGCGTGGGACATCCTCTCGCCGCAGTTTATGTACGGCTCGAAGG
CGTACGTGAAGCACCCGGCGGACATTCCGGACTACCTGAAGCTCTCGTTCCCGGAGG
GCTTTAAGTGGGAGCGCGTGATGAACTTTGAGGACGGCGGCGTGGTGACGGTGACG
CAGGACTCGTCGCTGCAGGACGGCGAGTTCATCTACAAGGTGAAGCTGCGCGGCAC
GAACTTTCCCTCGGACGGCCCGGTGATGCAGAAGAAGACGATGGGCTGGGAGGCGT
CGTCGGAGCGCATGTACCCCGAGGACGGCGCGCTGAAGGGCGAGATTAAGCAGCGC
CTGAAGCTCAAGGACGGTGGCCACTACGACGCGGAGGTGAAGACGACGTACAAGGC
GAAGAAGCCCGTGCAGCTGCCCGGCGCGTACAACGTGAACATCAAGCTCGACATTAC
GTCGCACAACGAGGACTACACGATTGTGGAGCAGTACGAGCGCGCGGAGGGCCGCC
ACTCGACGGGCGGCATGGACGAGCTCTACAAG

MVSKGEEDNMAIIKEFMRFKVHMEGSVNGHEFEIEGEGEGRPYEGTQTAKLKVTKGGPL
PFAWDILSPQFMYGSKAYVKHPADIPDYLKLSFPEGFKWERVMNFEDGGVVTVTQDSSLQ
DGEFIYKVKLRGTNFPSDGPVMQKKTMGWEASSERMYPEDGALKGEIKQRLKLKDGGHY
DAEVKTTYKAKKPVQLPGAYNVNIKLDITSHNEDYTIVEQYERAEGRHSTGGMDELYK

**P2A**

GGCTCGGGCGCGACGAACTTCTCGCTCCTCAAGCAGGCGGGCGACGTGGAGGAGAACCCGGGCCCG

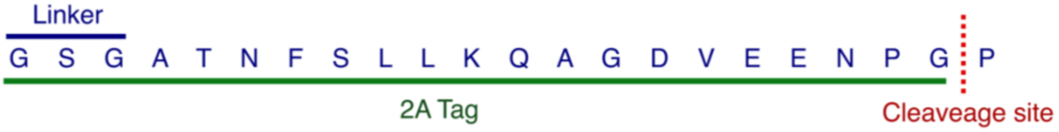


**Supplementary Figure 1.** Codon optimised DNA and protein sequence of mCherry fluorescent protein and porcine teschovirus-1 2A (P2A) pseudo-autolytic peptide.

*Malassezia furfur* CBS 14141 Chromosome 2, putative CDC25 Phosphatase (143 aa)

MSFSPPYQYMDADTLASELRRRTQAPNSIAVVDVRDDDYEGGHIVGAIHAPSGTFL
TRVDSLVGKELQDYERVVFHCSLSQQRGPKSARIYRETRDAAQAAGRVPTKEQAI
FVLRDGFAHFGPKFKNERDLVEDWDEEAWEWR*

*Malassezia furfur* CBS 14141 Chromosome 2, hypothetical Adiponectin Protein Receptor (632 aa)

MGSDVTPSESGSGVSTPPRVSCPSTPNHVHYTPVTAAALAALSDSIEGGPALRASQS
SRPPSISDHSEGTHTAAHSAVDAFSLPYWLAYLRSEAARHAHDIDQRMHAILESPD
SNDNVLKASISMMTHQLDVVYHALGNLSQRLPMGPSALYDSLPSQSDLSHKLQTL
MKDWEQHAHLPHSPLFPQTGPLSFGSLGWNASMEQDATASVPAWSYSSPFGLIPG
DWASMVHMPSREQVSDEVHRRLHAFVEQMHSMPARLSQAALPTSLAAWEAPAS
ELLHRVEEKLGEMSEQGRAGASHLVQRANQAVHDVEDVLYQAACELAREGRVLI
SYQSLPTLWRNNDCIHTGYRFIPVRNWTTLLGSIFQIHNETGNIHTHLSGLILVGALF
WFSGSLDSLTTTTDRWIQTLYLLAAAKCLVCSVSWHVMAGCADLNWFQCFACID
YTGISWLVAASLLTLVYNGFYCQPNLIAIYSVGVFLLGTTMGVLPWYPWFDDPKN
RTLRISLFVIMALVGLVPFTHGMYLHGFQHMVYFFSPIIPSIASYIAGVVVYALRFPE
KYWPGRFDLLGHSHQMWHIAIVLAIALHYRAILLFHKDRFTYSQVDGTCPSFADSL
PWSNAAVGGWWPQAGKALGAL*

**Supplementary Table 2.** Protein sequences of hypothetical genes located both upstream and downstream of the random mutagenesis insertion point in mf::mc-27 strain.
